# memerna: Sparse RNA Folding Including Coaxial Stacking

**DOI:** 10.1101/2023.08.04.551958

**Authors:** Eliot Courtney, Amitava Datta, David H. Mathews, Max Ward

## Abstract

Determining RNA secondary structure is a core problem in computational biology. Fast algorithms for predicting secondary structure are fundamental to this task. We describe a modified formulation of the Zuker-Stiegler algorithm with coaxial stacking, a stabilizing interaction in which the ends of multi-loops are stacked. In particular, optimal coaxial stacking is computed as part of the dynamic programming state, rather than inline. We introduce a new notion of sparsity, which we call *replaceability*. The modified formulation along with replaceability allows sparsification to be applied to coaxial stacking as well, which increases the speed of the algorithm. We implemented this algorithm in software we call *memerna*, which we show to have the fastest exact RNA folding implementation out of several popular RNA folding packages supporting coaxial stacking. We also introduce a new notation for secondary structure which includes coaxial stacking, terminal mismatches, and dangles (CTDs) information.

## 1 Background

New techniques have enabled an explosion in the number of known RNA sequences [1]. Many of these sequences are thousands on nucleotides long [2]. For example, SARS-CoV-2 is approximately 30 kilobases long [3]. Because these RNAs appear to be involved in important cellular functions [2], [4]–[6], there is interest in their structures, as structure typically determines function. In particular, determining the secondary structure of RNA is an important problem in RNA bioinformatics, as it largely determines RNA structure overall [7].

Techniques to determine RNA secondary structure can broadly be divided into computational and non-computational methods. Non-computational methods, such as nuclear magnetic resonance imaging and X-ray crystallography, can be very accurate. Since RNA is more labile than DNA, these techniques require expertise, time, and are expensive [8]. This can make such methods impractical in many cases. Thus, non-computational methods often form a gold standard by which to judge the more widely applicable computational methods.

There are different types of computationally based techniques. Comparison-based techniques use a set of homologous sequences, i.e. sequences with a common ancestor that perform the same function and should therefore have conserved structure, to find a common structure. In contrast to these, *de novo* techniques attempt to predict RNA structure from just the sequence. We shall refer to both these methods as *RNA folding* algorithms. Comparison-based folding algorithms are generally more accurate. These either require many homologous RNAs [9] and can be intractably slow to solve optimally for long RNAs [10]–[15] or use heuristics which may not produce the optimal solution or any solution [16]. Additionally, folding algorithms that perform comparison based structure prediction typically must solve the *de novo* folding problem too, as they build upon the same ideas.

Modern RNA folding algorithms are generally based on recursions first proposed by Zuker & Stiegler [17]. These recursions have grown to include the nearest neighbor model of RNA free energy change. This model is a gold standard and was developed by Turner *et al*. [18]–[23]. Popular modern RNA folding software packages include an implementation of a Zuker-Stiegler based algorithm using some subset or near-subset of the nearest neighbor thermodynamic model [24]–[27]. While the Zuker-Stiegler algorithm has proved to be a robust and useful tool, it requires *O*(*N* ^3^) computation time for computing the minimum free energy structure (with no pseudoknots), where *N* is the length of the RNA sequence. This requirement becomes intractable for longer RNAs.

Techniques have been developed to speed up the Zuker-Stiegler algorithm and simpler variants. These have generally been focused on applying variants of the Four-Russians speedup, or on sparsification, and often make simplifying assumptions about the thermodynamic model. Recently, LinearFold [28], [29] was introduced, which uses a beam search to speed up folding to *O*(*N*) and is much faster than existing RNA folding packages, although does not necessarily compute the exact solution given the energy model.

The Four-Russians speedup typically involves partitioning a matrix into square blocks, doing pre-computation on these, and achieving a speedup with the right block size and matrix values. Variants of this technique have been applied to RNA folding [30]–[32]. However, to our knowledge, all of these assume a simplified view of RNA thermodynamics, similar to the earlier algorithm of Nussinov & Jacobson [33].

Other approaches to speeding up RNA folding include *sparse folding*. These techniques modify the Zuker-Stiegler algorithm by not considering configurations that could not be in the optimal solution. Examples include the work of Backofen *et al*. [34], and Will & Jabbari [35]. These approaches speed up folding (empirically close to *O*(*N* ^2^)) while still computing the exact optimal structure. Further, they support most of the complete nearest neighbor model. However, they do not include the sparsification of coaxial stacking or dangling ends, which are motifs that are known to stabilize RNA structures [36].

We present a new sparse folding algorithm that includes the full nearest neighbor model including coaxial stacking, terminal mismatches, and dangling ends (CTDs). We also introduce a new conceptual framework that we use to enable sparse folding with CTDs, as well as to better understand sparse folding. Lastly, we introduce a new notation for RNA secondary structure that includes CTD information.

This algorithm is implemented by our RNA folding package, memerna [37], which is available at https://github.com/Edgeworth/memerna/tree/release/0.1. Results show that our algorithm is the fastest complete and exact (supporting the largest subset of the nearest neighbor energy model) RNA folding algorithm available at time of writing.

## 2 Energy Model

The nearest neighbor model for predicting the free energy of RNA structures is quite detailed. It is the result of many optical melting experiments, and years of analysis [18]–[22], [38]–[42]. We give a partially abstracted overview of this model.

The thermodynamic model is called the nearest neighbor model because it computes energy by decomposing structures into localized regions called *loops*. Formally, Zuker and Sankoff define a loop, closed by a base pair (*ost, oen*), to be the set of *accessible* bases from (*ost, oen*). A base *k* is accessible from (*ost, oen*) if, for all other base pairs (*ist, ien*), it is not the case that *ost < ist < k < ien < oen* [17]. That is, there are not any base pairs “between” (*ost, oen*) and *k*. There could be accessible bases from (*ost, oen*) that are in base pairs—those base pairs are called interior base pairs to the closing base pair (*ost, oen*). The names *ost, oen, ist*, and *ien* stand for “outer start”, “outer end”, “inner start”, and “inner end” respectively.

We will now describe the different types of motifs used to compute the free energy of a secondary structure. We also introduce some notation—the *dot-bracket* notation [43]. Since we assume there are no pseudoknots (a pseudoknot is a set of pairs (*i, j*) and (*k, l*) with *i < k < j < l*), we can write the base pairs of an RNA using nested parentheses. Two bases are in a base pair if their parentheses match, and not bonded if they have a dot instead. For example, in the primary structure “GAAAC”, the dot-bracket string “`(…)`” indicates that the G and C bases are paired, and the rest are not. We can then write the full secondary structure like this:

~~~
                          GAAAC
                         (…)
~~~

This notation is useful for visualizing the structure of loops when thinking about dynamic programming algorithms. We will also use angle brackets (“`<…>`“) to represent any substructure—i.e. any valid sequence of parentheses and dots.

### 2.1 Helices or “Stacking”

Helices are contiguous lengths of base pairs. To couch it in the definition of loop we provided, they are loops with exactly one interior base pair, (*ist, ien*) to the exterior base pair (*ost, oen*), and no unpaired bases. That is, *ist* = *ost*+1 and *ien* = *oen* − 1 [44]. These are energetically favorable because of what are called stacking interactions—the planes of adjacent base pairs are stabilized by being adjacent and parallel. In the Turner 1999 and 2004 models, these helices are assigned stabilizing energies based on which base pairs are adjacent. For example, if an AU pair is followed by a GC pair, it is assigned a free energy of -2.08 kcal/mol. There is also a case where one special sequence of four base pairs is given a different free energy, but as far as we can tell this is not implemented by any RNA folding packages [24], [25], including memerna. This is now known to be an artifact, as shown by updated nearest neighbor rules [45].

Whenever a helix is ended or started by a base pair that is AU, GU, UG, or UA, a special *AU/GU penalty* is applied. In the case of a length one helix it is applied twice—once when it starts and once when it ends.

In dot-bracket notation, helices are shown as concentric parentheses, for example:

~~~
                          ((((<…>))))
~~~

We have summarized the computation in Algorithm 1.

### 2.2 Hairpin Loops

Having helices necessarily means that sometimes we must form loops. These are generally energetically unfavorable. A hairpin loop is a sequence of at least three unpaired bases closed by a base pair—that is, it is a loop with no interior base pairs. There has been a lot of research done on the folding stability of hairpin loops, and there are many special interactions that have been observed. For example, there are special tetraloops (a hairpin loop with four unpaired bases in it) that have their own recorded energy values in the Turner 2004 and Turner 1999 models [18], [19].

In dot-bracket notation, hairpin loops look like one pair of parentheses, with at least three dots in the middle, for example:

~~~
                          (…)
~~~

Hairpin loop energy computation is described in Algorithm 2. Constants (like “GGFirstMismatch”), are recorded in the Nearest Neighbour Database (NNDB) [40].

### 2.3 Bulge Loops

Bulge loops are loops that have exactly one interior base pair, (*ist, ien*), and have no unpaired bases on one side of the exterior (closing) base pair, (*ost, oen*). That is, exactly one of *ist* = *ost* + 1 or *ien* = *oen* − 1 is true. This is because the “bulging” unpaired base is pushed out and the stacking interactions can continue as if it were not there between the otherwise adjacent base pairs. Like hairpin loops, Turner 2004 defines a complicated piecewise function that depends on how many unpaired bases there are and what kind of bases are inside the bulge loop [40].

In dot-bracket notation, bulge loops have two parentheses adjacent to each other on one side, and at least one unpaired base between the two parentheses on the other side, for example:

~~~
                          ((<…>)..)
~~~

See Algorithm 3 for the computation.

### 2.4 Internal Loops

Internal loops look like bulge loops, but they also have one or more unpaired bases on the other side. Formally, *ist > ost* + 1 and *ien < oen* − 1. These are often lumped together with bulge loops and stacking, as they can be generalized to something sometimes referred to as a *two-loop* [44], since the only difference between them is that internal loops have unpaired bases on both sides. There are, again, special cases for internal loops up to size two by three (two unpaired bases on one side, three unpaired bases on the other).

In dot-bracket notation, internal loops look like bulge loops, except they must have at least one unpaired base between the two parentheses on both sides, for example:

~~~
                          (…(<…>)..)
~~~

Algorithm 4 shows the algorithm for internal loops, and Algorithm 5 shows the combined algorithm for two-loops.

### 2.5 Multi-loops

Multi-loops are loops with multiple interior base pairs. The areas closed by the base pairs in a multi-loop are called branches, including the closing (or outer) base pair. Some additional interactions have been added since the original Zuker-Stiegler algorithm, such as *coaxial stacking*, which is where adjacent branches can have stacking interactions between their closing base pairs.

Multi-loops are poorly predicted by the Turner models. Even though the Turner 1999 and 2004 models give logarithmic and asymmetry based equations describing the free energy of multi-loops, in practice, these are not used for free energy minimization—instead, an affine approximation is used [46]–[51].

Memerna uses the same affine approximation as RNAstructure, which is where:

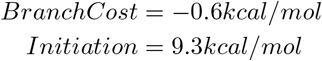

But, there is another setting commonly used, where *BranchCost* = −0.9*kcal/mol* [19].

In dot-bracket notation, a multi-loop looks like an outer pair with at least two branches inside, separated by any number of unpaired bases. Here is an example of one:

~~~
                          (..(<…>)(<…>).)
~~~

See Algorithm 6 for the implementation.

### 2.6 Coaxial Stacking, Dangling Ends, and Terminal Mismatches

*Coaxial stacking* is a stacking interaction between the closing base pairs of adjacent branches, or branches separated by one unpaired base, and has been shown to increase the accuracy of RNA secondary structure prediction [36], [40]. Coaxial stacking is one of the three possible interactions that can occur with branches in the Turner 2004 model, the other two being dangling ends and terminal mismatches. A *dangling end* is the interaction of an unpaired base adjacent to a branch. There are two types, 3^*′*^ and 5^*′*^, depending on which side of the branch it is on—if it is on the 5^*′*^ side of the branch, it is called a 5^*′*^ dangle. If a branch has an unpaired base adjacent on both sides, they cannot both dangle. Instead, they can form what is called a *terminal mismatch*, which is counted differently in the energy model. A branch or unpaired base can only be involved in at most one of these three interactions, and to compute the MFE structure, we need to take the combination of interactions that gives the lowest free energy. This significantly complicates the Zuker-Stiegler prediction algorithm, and existing sparsification techniques do not handle it [35].

There are two types of coaxial stacking: flush coaxial stacking, and mismatch-mediated coaxial stacking. In flush coaxial stacking, two branches directly adjacent to each other stack on top of each other in an energetically favorable way. It looks like this in dot-bracket notation: “`(<…>)(<. >)`”. Mismatch-mediated coaxial stacking is similar but has an unpaired base in-between the branches: “`L(<…>).(<. >)R`”, and is counted differently in the energy model. The unpaired base can form a terminal mismatch with either L or R in the previous example. The energy is determined by looking up a value based on the two bases in the terminal mismatch, and the two bases in the base pair of the branch straddled by the terminal mismatch.

### 2.7 Coaxial Stacking, Dangling Ends, and Terminal Mismatches notation

We present an extension to the dot-bracket notation which also includes CTD information. Current dot-bracket notation has no way of representing the CTD information, even though popular RNA packages produce secondary structures with particular CTDs.

Parentheses are replaced by square brackets (to differentiate between a secondary structure with no CTDs and one with none specified). Dangles are represented by “5” and “3” for 5^*′*^ and 3^*′*^ dangles respectively. Terminal mismatches are represented by “m” and “M”, for the unpaired bases that are on the 5^*′*^ side and 3^*′*^ side, respectively. It is necessary to use two different symbols here, rather than lower-case “m” for both, as there would otherwise be ambiguous cases. The Turner 2004 model allows branches to be in two coaxial stacks: one as an internal branch to a multi-loop, and another as the outer branch of a multi-loop. In the following example, branches are indicated by the letter they contain, except for the outer branch of the multi-loop on the right, which is B. The branch B can be involved in a flush coaxial stack with both A, and C or D.

~~~
                          (.A.)((.C.)(.D.))
~~~

To be able to store both of these interactions, we put the interaction of the branch as an internal branch in the location where the left square bracket would be, and the interaction of the branch as the outer branch of a multi-loop where the right square bracket would be. A branch in a coaxial stack can either interact with the previous branch (in the 5^*′*^ direction), or the next branch (in the 3^*′*^ direction), which is encoded as “p” for previous, and “n” for next. In the case of an outer branch, these are capitalized, and next refers to the first internal branch in that multi-loop.

Similarly, an outer branch can also be involved in terminal mismatches or dangles on both sides of the branch. For its interaction inwards, it is a 180 degree rotation so dangles and mismatches are apparently “swapped”. For example:

~~~
                          m[M[…]..[.]m]M
                          .[3[…]..[…].]3
~~~

Further, in the example shown in Figure 1, the first line indicates the names given to the branches below, which are in normal dot-bracket notation. The last line shows the CTD interactions. Branch A is involved in a terminal mismatch, and a 3^*′*^ dangle. Branches B and C have no interactions. Branch D is in a flush coaxial stack with E. Branch F has a 5^*′*^ dangle. Branch E additionally, as an outer loop, is involved in a flush coaxial stack with G.

**Figure 1:**
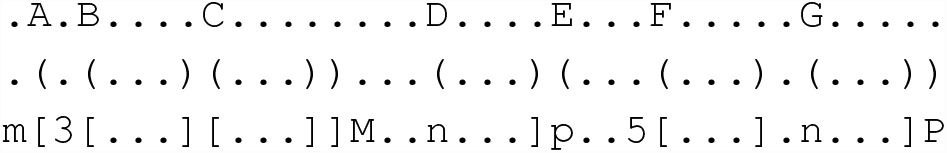
CTD notation

## 3 Sparse folding with coaxial stacking

### 3.1 Reformulating coaxial stacking

Adding coaxial stacking, terminal mismatches, and dangles (henceforth referred to as *CTDs*) to the Zuker-Stiegler algorithm significantly complicates it, although the time complexity does not change. In order to apply sparse folding, we modify the traditional recursions for the Zuker-Stiegler algorithm to represent sufficient coaxial stacking information in the dynamic programming state.

In particular, our formulation considers branches separately when finding the optimal CTD configuration, whereas normally coaxial stacks are optimized by trying every possible *split point* explicitly, in a loop. A split point (also called a *pivot point*) here is an index which has a branch both to the left and right of it. For a particular split point, the existing algorithm looks at the branch to the left and the right and compute the free energy contribution of those two branches forming a coaxial stack.

The cases for each kind of CTD configuration are shown in the following tables. Each case describes a potential arrangement of branches in a multiloop (or exterior loop). Table 1 shows the cases used to compute the exterior loop and Unpaired table. Table 2 shows the cases used to compute the Paired table.

**Table 1:** Unpaired and external function cases.

| Case | Shape | Description |
| --- | --- | --- |
| NoCTD | [<...>]<...> | One branch with no CTDs |
| Dangle3' | [<...>]3<...> | One branch with a 3' dangle |
| Dangle5' | 5 [<...>]<...> | One branch with a 5' dangle |
| Mismatch | m [<...>]M<...> | One branch with a terminal mismatch |
| LeftCoax | mn<...>]Mp<...>]<...> | One branch that must be in a left mismatch mediated coaxial stack with something to the right of it |
| RightCoaxFwd | n<...>]mp<...>]M<...> | One branch that must be in a right mismatch mediated coaxial stack with something to the right of it |
| FlushCoaxWC | n<...>]p<...>]<...> | One branch that must be in a flush coaxial stack with something to the right that is closed by a Watson-Crick pair |
| FlushCoaxGU | n<...>]p<...>]<...> | One branch that must be in a flush coaxial stack with something to the right that is closed by a GU pair |

**Table 2:** Paired function cases.

| Case | Shape | Description |
| --- | --- | --- |
| Multiloop | [<...><...>] | Any valid multiloop contents |
| OuterDangle3' | [3<...><...>] | Outer loop forms a 3' dangle |
| OuterDangle5' | [<...><...>5] | Outer loop forms a 5' dangle |
| OuterMismatch | [M<...><...>m] | Outer loop forms a mismatch |
| LeftOuterCoax | [Mp<...>]<...>mN | Outer loop forms an outer mismatch mediated coaxial stack with a branch on the left |
| RightOuterCoax | [M<...>n<...>]mP | Outer loop forms an outer mismatch mediated coaxial stack with a branch on the right |
| LeftInnerCoax | [mp<...>]M<...>N | Outer loop forms an inner mismatch mediated coaxial stack with a branch on the left |
| RightInnerCoax | [<...>mn... ]MP | Outer loop forms an inner mismatch mediated coaxial stack with a branch on the right |
| LeftFlushCoax | [p<...>]<...>N | Outer loop forms a flush coaxial stack with a branch on the left |
| RightFlushCoax | [<...>n<...>]P | Outer loop forms a flush coaxial stack with a branch on the right |

We break down flush coaxial stacking into two types: *inner*, and *outer*. The outer type happens when one of the branches is the outer loop in a multi-loop. We also break down mismatch mediated coaxial stacking into six types: *left, right, left outer, right outer, left inner*, and *right inner*. The first two do not involve an outer loop, the rest do.

### 3.2 Replaceability and monotonicity

Sparse folding is a way of reducing the number of split points considered when computing functions like Paired or Unpaired. It makes the time complexity *O*(*N* ^2^) in the average case, with some assumptions [52]. It is not a new technique [34], [35], [52]. But, it has not been applied to coaxial stacking yet – doing so requires entirely new recursions for coaxial stacking, which we have described in Appendix B.

In existing literature, sparse folding hinges upon the concept of monotonicity. Suppose we are computing *Unpaired*(*st, en*) and are considering placing a branch at (st, piv) — that is:

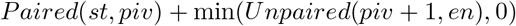

If we previously found some *Paired*(*st, lessthanpiv*) *< Paired*(*st, piv*) where *lessthanpiv < piv*, because of the triangle inequality of Unpaired, we know that choosing the split point at *lessthanpiv* must be at least as good as choosing it at *piv*. This lets us then build a candidate list *C*(*st, en*) of split points for each *Unpaired*(*st, en*), which must be monotonically decreasing in *Paired*(*st, p*) for all *p* in *C*(*st, en*). The candidate list is used to reduce the number of candidate split points we need to search over for future computation.

The existing sparse folding technique, using monotonicity, is not directly applicable to coaxial stacking. We need to distinguish monotonicity from the notion of *replaceability*. In the previous example, we saw that we did not need to consider splitting at *piv* because the split point at *lessthanpiv* was at least as good. This carries with it the idea that we could replace *Paired*(*st, piv*) with *Paired*(*st, lessthanpiv*) and still get a valid structure. We say that *Paired*(*st, piv*) is *replaceable* by *Paired*(*st, lessthanpiv*). These two ideas of replaceability and monotonicity may not have been separated much in existing work. Thinking of them as separate things, with monotonicity as an additional optimization to replaceability is necessary to apply them to coaxial stacking.

Suppose we are trying to compute the value of *Unpaired*(*st, en*), at some split point *piv*. We check if *Paired*(*st, piv*) + min(*Unpaired*(*piv* + 1, *en*), 0)) is better than any split point we have checked so far. But, the value of *Unpaired*(*st, piv*) (ignoring the case where *Unpaired*(*st, en*) tries pairing at (*st, en*)) can tell us if we could do better than *Paired*(*st, piv*). That is, if we could *replace Paired*(*st, piv*) with a different structure that has lower energy without making our function mean something else, then we do not have to consider the split point *piv*. Note that it does not change the meaning of our function if the best structure *Unpaired*(*st, piv*) gives us does not start with a branch at *st*. That is handled when we check the case where *Unpaired*(*st, en*) = *Unpaired*(*st* + 1, *en*).

Now we have a condition for when we do not need to check some split point *piv*: *Paired*(*st, piv*) *> Unpaired*(*st, piv*) (or ≥ if we can check *Unpaired*(*st, piv*) before it considers the *Paired*(*st, piv*) case). From this, we can build up *candidate lists*, which are lists of split points we need to consider for each start point *st*. If we implement the Zuker-Stiegler algorithm iteratively, with *st* in the outer loop and *en* in the first inner loop, then we only need one candidate list. Every iteration of the outer loop we clear it, and build up the list of split points using each *Paired*(*st, en*) we look at by recording the *en* index as a split point *piv* that we must check in later iterations. When we are computing some *Unpaired*(*st, en*), we will have already considered putting each possible split point *st < piv < en* into the candidate list.

Another observation is that these candidate lists can be strictly monotonically decreasing. We know that *Unpaired*(*st, en*) ≤ *Unpaired*(*st* + *k, en*), because checking *Unpaired*(*st, en*) = *Unpaired*(*st* + 1, *en*) is one of the cases. This corresponds to the idea that we could always pad out *Unpaired*(*st* + *k, en*) with unpaired bases (that cost zero in the Turner 2004 model) until it spans from *st* to *en*. This means that there is no point in considering some split point *piv* if there was an earlier split point *asgoodaspiv < piv* that was just as good—we could just pad out *Unpaired*(*piv* + 1, *en*) to *Unpaired*(*asgoodaspiv* + 1, *en*) and the free energy would not change.

Let us work out a few conditions for a structure to be replaceable in the context of one of our functions. Firstly, the structure with which we consider replacing it with needs to be representable and considered by that function. Secondly, we need to know the exact free energy contribution of the structure we are considering replacing. For example, if we are trying to place a coaxial stack, replacing the left branch with *Unpaired* will destroy that coaxial stack. The energy from the left branch and that of the coaxial stack itself is the “gain” we get from not replacing it, so we need to know the total free energy of these contributions.

One of the existing recurrence relations (for example, this used in RNAstructure) for CTDs works by computing a table that stores the best pair of branches and CTD configuration for some range (*st, en*). The first branch in the pair must start at *st*, and the second must end at *en*. This table is computable in *O*(*N* ^3^) time, by iterating over every possible splitting point. This works because two branches involved in a coaxial stack have to be adjacent, or only have one intervening base. This table is used for placing coaxially stacked structures inside a multi-loop, but interactions with the outer loop need to be considered separately. If there is an outer loop at (*st, en*), it tries all possible branches that start at *st* + 1 and *st* + 2 (for mismatch mediated, and flush coaxial stacks respectively), and all possible branches that end at *en* − 2 and *en* − 1. There are only approximately 2*N* of these at most, so the overall complexity is still *O*(*N* ^3^) for the case of an outer loop as well.

This recurrence relation does not lend itself to sparse folding, however. The replaceability idea does not hold: if we replaced the left branch with *Unpaired*(*st, en*), the meaning of the table would change to be “the best structure with at least two branches, one ending at *en*, possibly with coaxial stacking”. But, this is not representable with the current recurrence relation, so it would give wrong results. Even if we fixed all these issues, it would not be *canonical*—it would consider the same structure multiple times, so e.g. exhaustive suboptimal folding [53] would not work properly—and would have worse sparsity properties than our formulation.

To do sparse folding with coaxial stacking, instead of having one candidate list, we use fourteen—one for each interaction type—as shown in Table 3 and defined below. Fewer lists would also work, but it would need to consider each potential interaction at every split point in the candidate list, only one of which is potentially worth looking at. Also, it makes it easier to pre-compute the energy contributions, rather than re-computing them in every split loop.

**Table 3:**
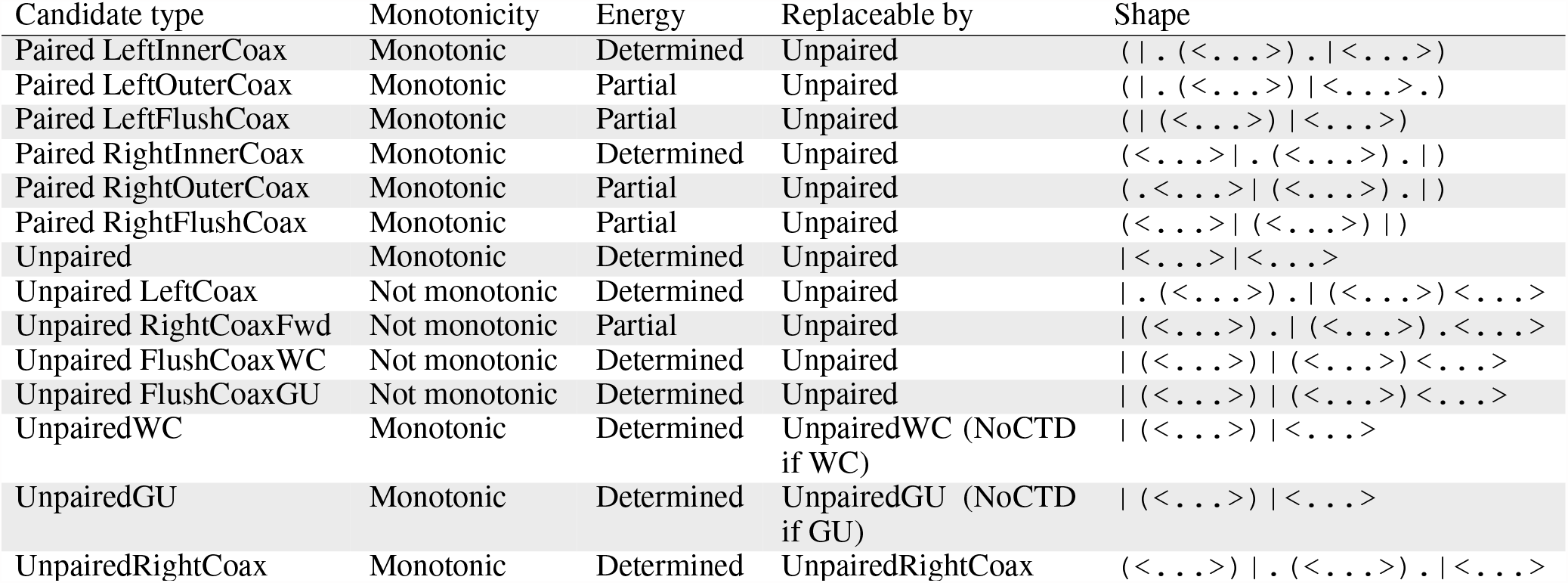
Candidate list types.

The idea of a replaceable structure changes slightly now too, and we will call it a *block*. Before, it was just a branch like this: “`(<…>)`”. Now, we define it as a block of affected area. For example, the block (*st, en*) containing a 3^*′*^ dangle is the whole sub-structure “`(<…>).`”. The energy of that block is the energy of everything it contains or references, except from parts that we have no way of knowing the energy contribution of.

Table 3 shows the fourteen different candidate list types, whether they can be monotonic or not, whether we fully know the energy contribution from each block at the time of considering it, what they are replaceable by, and the block that is replaceable in their shape (denoted with |…|). The non-monotonic ones are non-monotonic because, for the right split in their computation, they do not add the value of a function like *Unpaired* which obeys the triangle inequality [52] or *f* (*st, en*) ≤ *f* (*st* + *k, en*).

For example, Unpaired RightCoaxFwd is quite a bad candidate list, because it cannot prune many split points. It looks at making a right coaxial stack in the *Unpaired* table with the left branch (*st, en* − 1) in the block (*st, en*), and the best right branch from *UnpairedRightCoax*—the case itself looks like this: *Paired*(*st, en* − 1) + *UnpairedRightCoax*(*en* + 1, …). It does not have monotonicity, and we only know the energy of the left branch (*Paired*(*st, en* − 1)), making the energy only partially determined. In these cases, we can assume that it has the lowest possible free energy contribution from a mismatch mediated coaxial stack. This means we will put unnecessary candidates into the list, but the alternative is making eight extra candidate lists: one for each combination of right terminal mismatch base, and right branch base pair (either Watson-Crick or GU/UG). Assuming the worst case for free energy contribution lets us maintain replaceability because we could place something in the area from *st* to *en* that would make up for cost of not placing the coaxial stack.

For each case or block, we need to find what it is replaceable by. Usually, this is *Unpaired*. But, the *UnpairedRightCoax*(*st, en*) function must start with a branch at *st*, since it gives the value of the best right half of a right coaxial stack. It can only be replaced by itself. The *UnpairedWC* and *UnpairedGU* functions are similar—the branch starting at *st* must be closed by a Watson-Crick or GU/UG pair respectively, so they also can only be replaced by themselves. Since these functions are monotonic, it does not matter if we do not replace them with something else, because we still get the speed-up from monotonicity.

## 4 Empirical performance

### 4.1 Methodology

We have analyzed seven different configurations of packages, described in Table 4.

**Table 4:** Program configuration descriptions.

|  |  |
| --- | --- |
| LinearFold | LinearFold program |
| RNAstructure | RNAstructure Fold program |
| SparseMFEFold | SparseMFEFold program |
| ViennaRNA-d2-noLP | ViennaRNA with d2 option and no lonely pairs |
| ViennaRNA-d3 | ViennaRNA with d3 option |
| ViennaRNA-d3-noLP | ViennaRNA with d3 option and no lonely pairs |
| memerna | memerna fold program |

SparseMFEFold is an RNA folding package that does sparse folding including sparse memory optimizations [35], but it does not handle CTDs at all. ViennaRNA-d2 is ViennaRNA [24] with the `-d2` option, where it assumes dangles everywhere (and double counts them). ViennaRNA-d3 has the `-d3` option, which makes ViennaRNA compute CTDs, although it does not handle coaxial stacking of the two inner branches in a three branch multi-loop [54]. LinearFold is a linear time MFE folding package using beam search [28], and we run it with the same semantics as `-d2`. RNAstructure is the Fold program of RNAstructure [25], which fully handles CTDs.

For MFE calculation, we measured two quantities: runtime and maximum memory usage. We executed each program five times for each datum and took the mean. For runtime, we measured the wall time. For the memory usage, we took the maximum resident set size over the whole execution of the program. We used GNU time with the following command to get these values: `/usr/bin/time -f %e %U %S %M`.

We used Python to conduct the benchmarking and analysis, using the `NumPy`, `Pandas`, and `matplotlib` libraries from SciPy [55], the graphing library `Seaborn` [56], and the statistical analysis library `statsmodels` [57]. The command lines used to execute each program are shown in Table 5. We ran the benchmarks on a machine with a Ryzen 5950X processor, 64 GB of RAM, running Arch Linux with gcc 12.2.1 and clang 15.0.7.

**Table 5:**
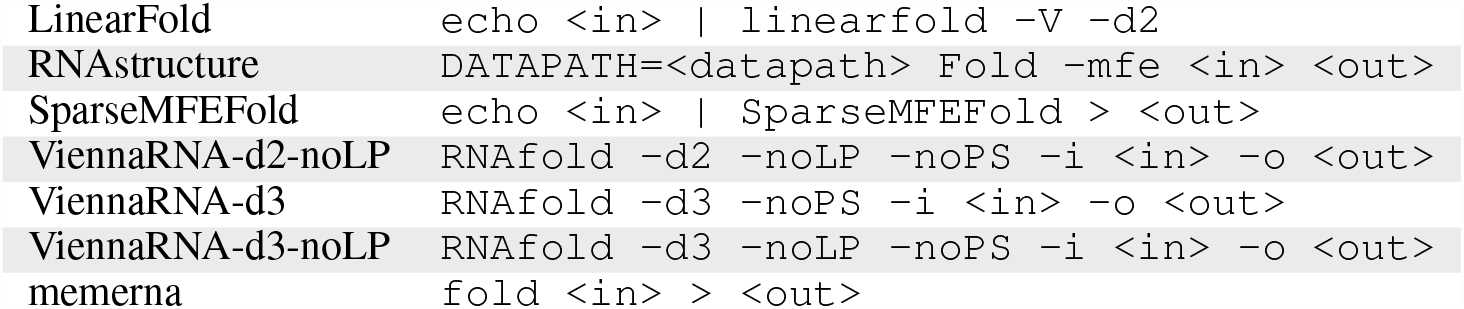
MFE run commands.

We generated a set of uniformly random sequences with lengths from fifty to three thousand, at intervals of fifty.

### 4.2 Results

The runtimes of seven configurations are displayed in Figure 2a, for each length of RNA in the random dataset. What we can see from this figure is that, in general, RNAstructure is the slowest, followed by ViennaRNA-d3, ViennaRNA-d2, SparseMFEFold, then memerna, which is the fastest exact algorithm, and then LinearFold (which is faster for lengths above 1800). Memerna does the same amount of “work” as RNAstructure (which handles CTDs completely) and more than ViennaRNA-d3, since ViennaRNA-d3 does not handle certain types of coaxial stacks [54]. ViennaRNA-d2 and LinearFold assume that every unpaired adjacent base dangles, and SparseMFEFold has the same behavior as ViennaRNA with the d0 option, which does not consider CTDs at all [35].

**Figure 2:**
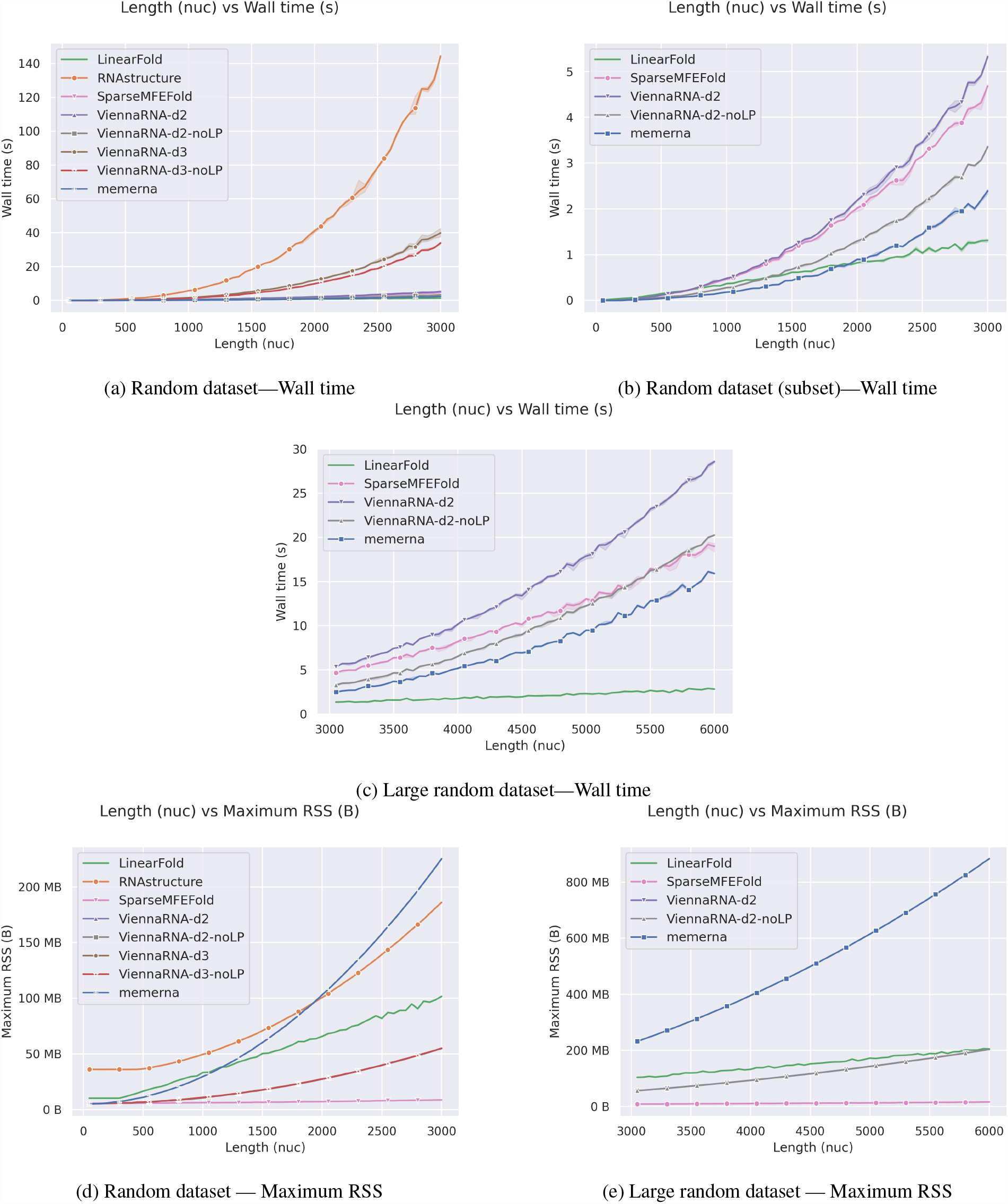
Wall time and memory usage

In Figure 2d we can see the memory usage, in terms of maximum resident set size. Overall, memerna had the highest memory usage, followed by RNAstructure, ViennaRNA, and finally SparseMFEFold, which unsurprisingly had the least memory usage. The higher memory usage of memerna is expected due to needing to store the candidate lists — it is still quite low and it is possible to run, for example, a 30kb sequence in a normal desktop computer.

We conclude from these figures that memerna is faster for exact folding than the tested packages for RNAs up to length six thousand, despite fully implementing coaxial stacking, although it uses more memory. It is faster than LinearFold up until RNAs of length 1800. It is possible to halve memerna’s memory usage by changing the free energy data type to `int16_t` (like RNAstructure) from `int32_t`.

## 5 Limitations

Our package memerna implements the Zuker-Stiegler algorithm with coaxial stacking, with some assumptions about the energy model. It assumes a Turner 04-like energy model, and is not as flexible as packages like RNAstructure or ViennaRNA as to what energy models can be specified. In particular, the unpaired base cost is not configurable from the Turner 04 model value of zero. These are not inherent limitations. Memerna could be modified to support a similar set of data table parameters with no change in speed, except for the unpaired base cost, which, if positive, would impact performance.

This formulation of the algorithm also depends upon the set of bases used — adding a new base type is not as easy as it is in RNAstructure [58], for example. This is because there are specific dynamic programming tables that encode what closing base pair is used for a branch. It is possible to modify the recursions to not depend on this, but it will reduce the sparsification since it would have to assume the minimum energy closing base pair to maintain replaceability.

## 6 Conclusion

By modifying the commonly used recurrences to include information required for coaxial stacking in the state, we were able to fully sparsify the Zuker-Stiegler algorithm with coaxial stacking. Empirically, this sped up folding significantly. Our implementation is approximately as fast as ViennaRNA with the d2 option (maybe slightly faster), but has a model-accurate implementation of coaxial stacking, terminal mismatches, and dangling ends, like that used in RNAstructure.

This is useful for folding large structures, such as SARS-CoV-2 which is approximately 30 kilobases long. It is also useful for folding many smaller structures. Computing the dynamic programming tables is also used as an input to other algorithms, for example, suboptimal folding.

An ostensibly obvious target for future work is to apply this to suboptimal folding and the partition function. This is likely to be hard or not useful since those algorithms work on multiple secondary structures or the entire thermodynamic ensemble, and sparsification is about skipping states that are not in one *particular* structure. It may be easier to apply it to Sankoff simultaneous alignment.

## 7 Source code

See https://github.com/Edgeworth/memerna/tree/release/0.1.

## A Pseudocode for energy functions

### Algorithm 1 Computes the energy of a helix where *r* is a primary sequence, with (*ost, oen*) as an outer base pair, and (*ist, ien*) as an inner base pair

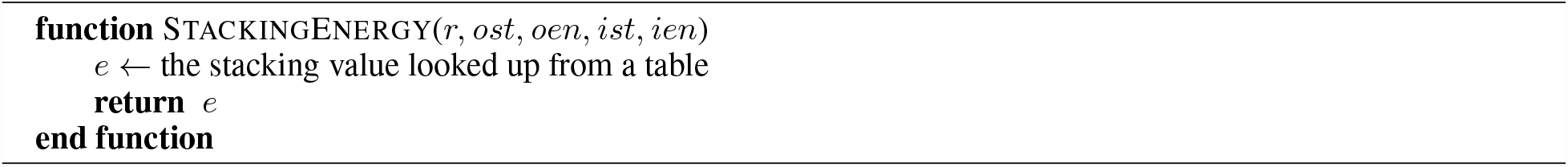

### Algorithm 2 Computes the energy of a hairpin loop, where *r* is a primary sequence, *r*[*st* : *en*] is the hairpin loop, and (*r*[*st*], *r*[*en*]) is the closing base pair

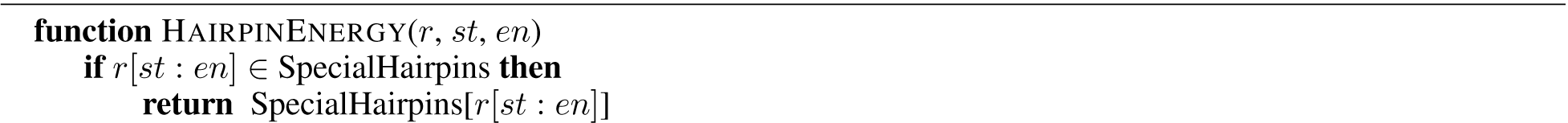
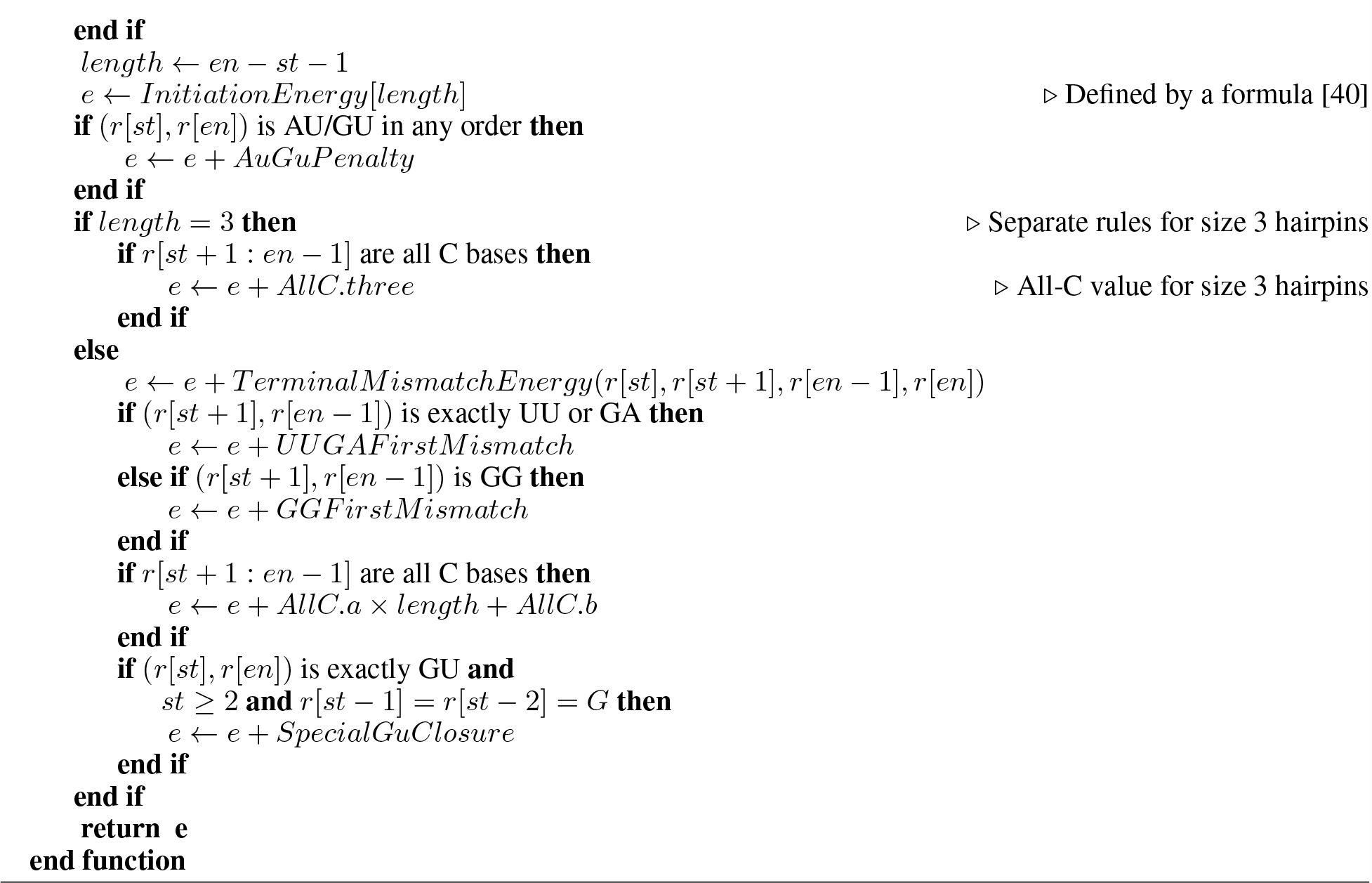

### Algorithm 3 Computes the energy of a bulge loop, where *r* is a primary sequence, (*ost, oen*) is the outer base pair, and (*ist, ien*) is the inner base pair

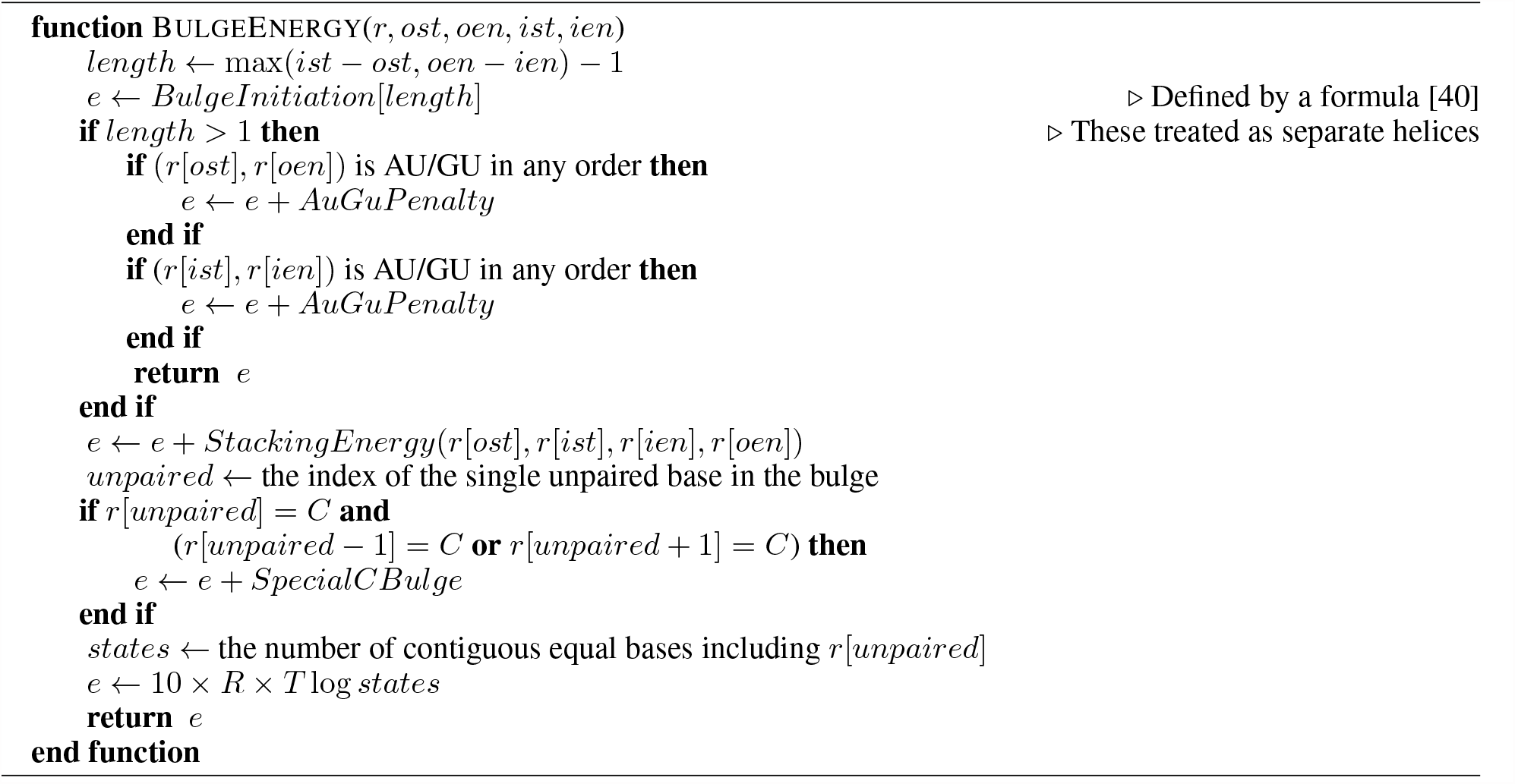

### Algorithm 4 Computes the energy of an internal loop, where *r* is a primary sequence, (*ost, oen*) is the outer base pair, and (*ist, ien*) is the inner base pair

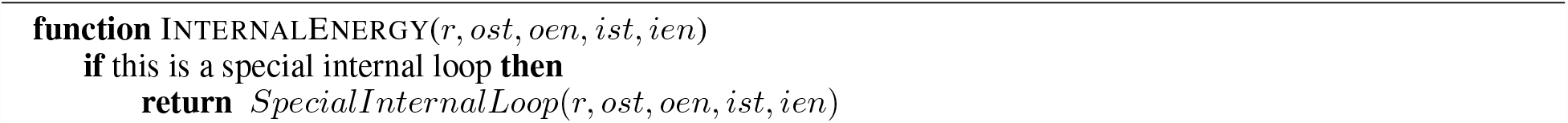
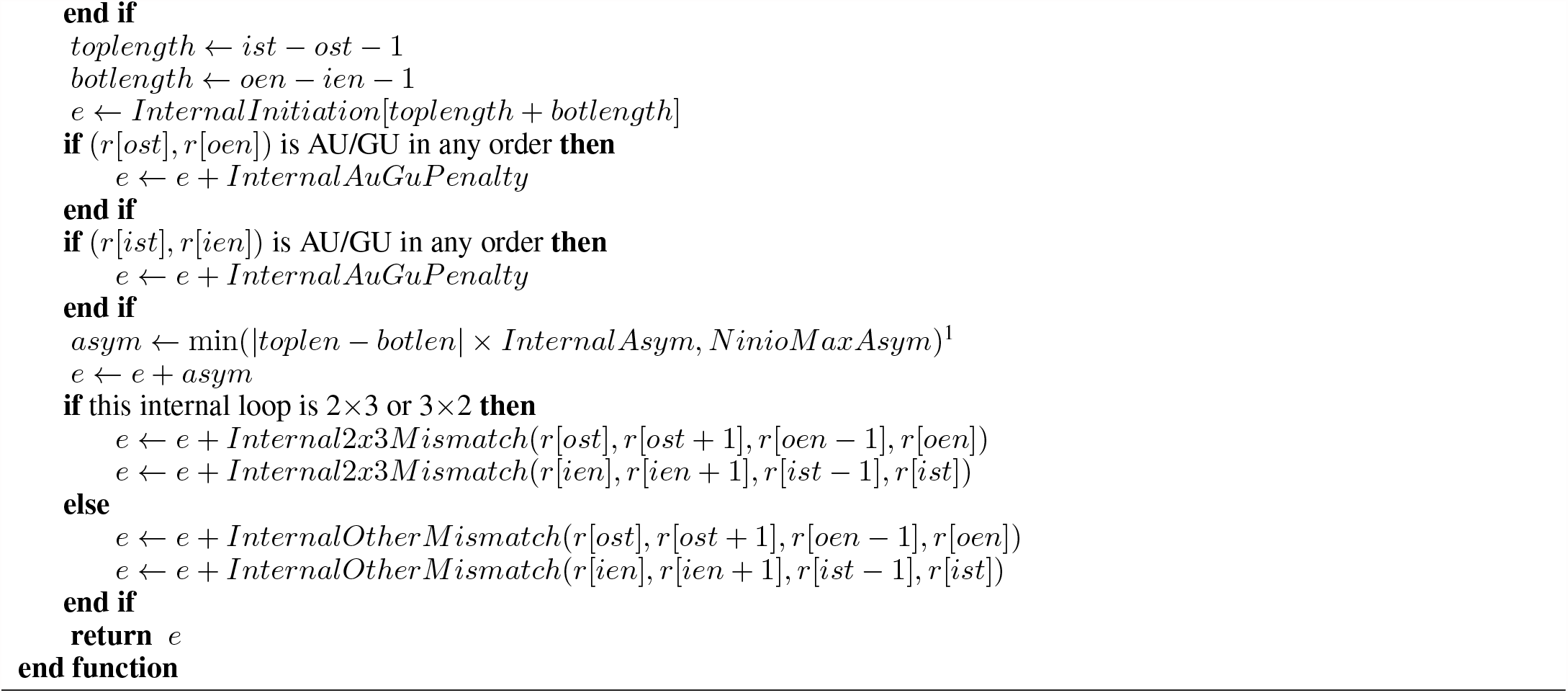

### Algorithm 5 Computes the energy of either an internal loop, a bulge loop, or stacking

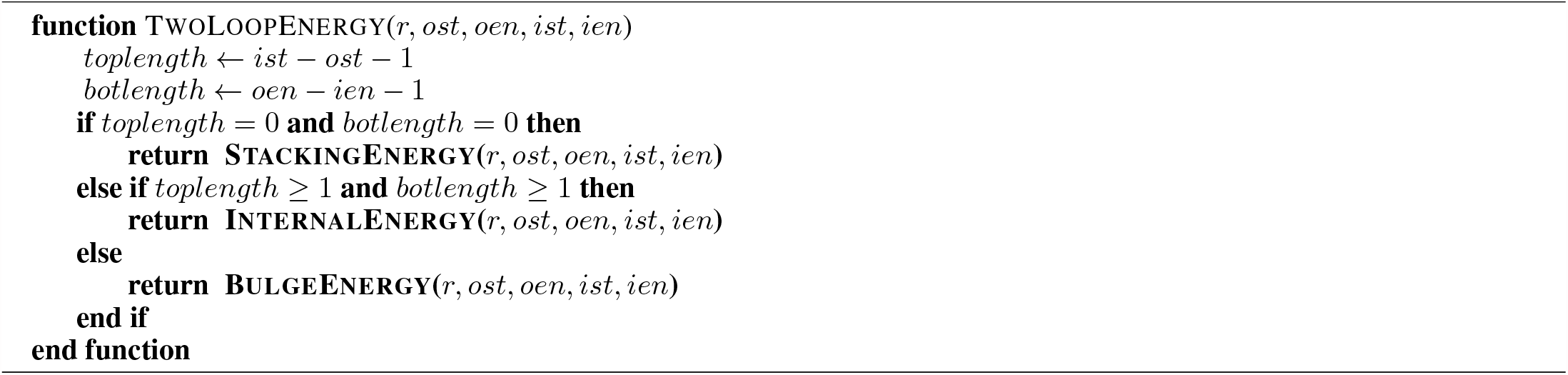

### Algorithm 6 Computes the energy of a multi-loop

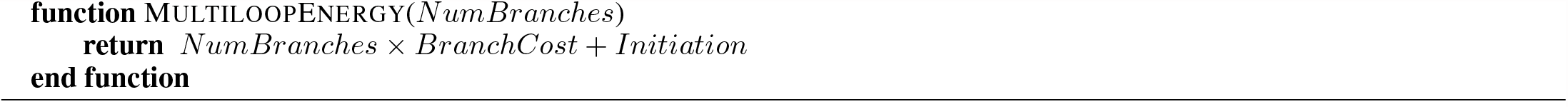

## B Dynamic programming recursions — details

This section contains the modified dynamic programming recursions in detail.

First, define the following helpers:

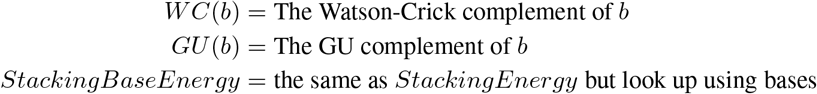

We start by again defining the function *Ext*, which computes the exterior loop:

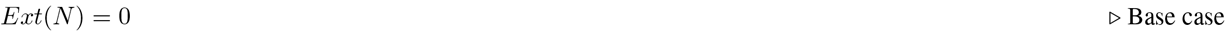

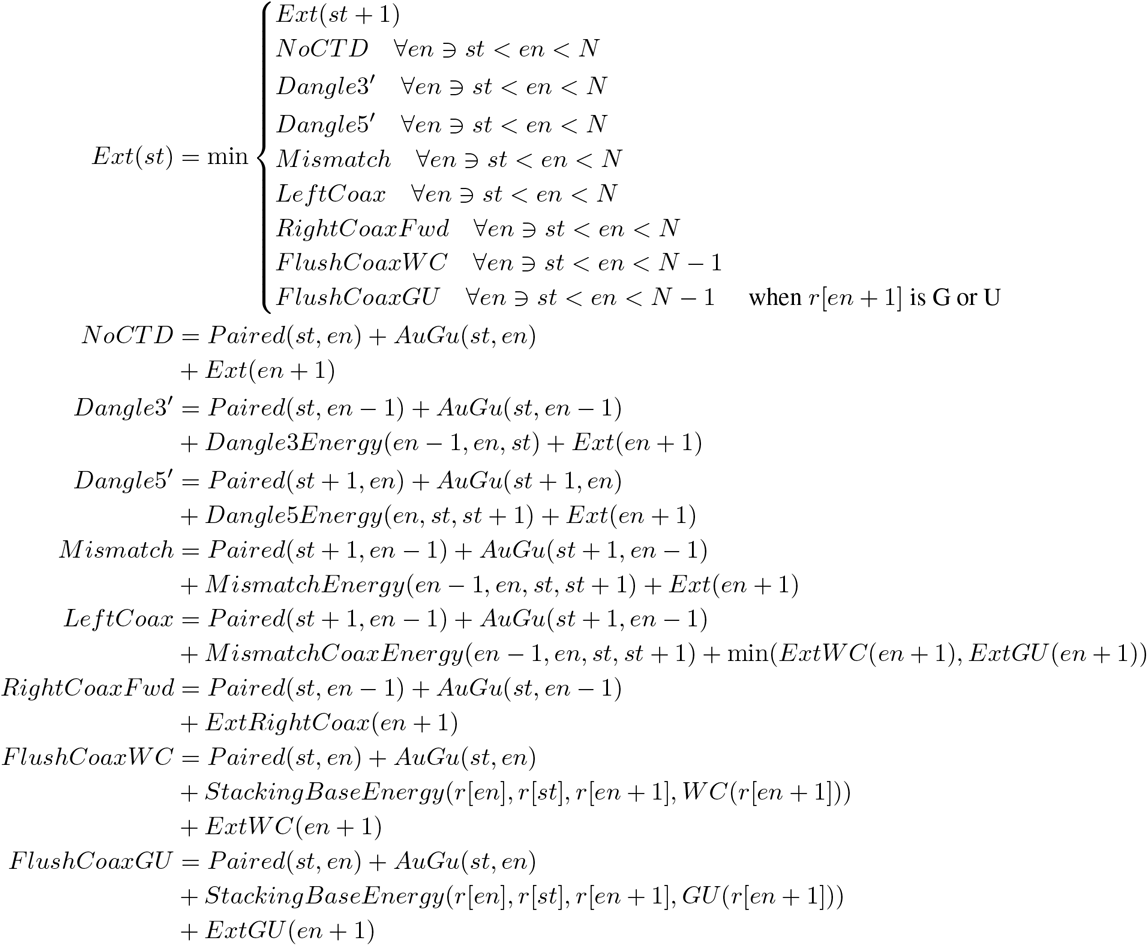

Table 1 shows a visual guide for each case.

For dealing with CTDs we need a few helper functions: *ExtGU, ExtWC*, and *ExtRightCoax*. The *ExtGU* (*st*) function returns the MFE given that *st* is in a GU or UG pair. The function *ExtWC* is similar, but for Watson-Crick pairs. Suppose we are trying to make a flush coaxial stack with the left branch at (*a, b*). We know the right branch starts at *b* + 1 and what base that is. If we knew whether it was closed by a Watson-Crick or GU/UG pair, we would know what the final base is, and be able to work out the energy. This is what these two functions are for. Left mismatch coaxial stacks also work with this method, since we trivially know all the bases. Right mismatch coaxial stacks are tricky and require the *ExtRightCoax*(*st*) function. It gives the MFE of the best structure given that a branch starts at *st* and is involved backwards in a right coaxial stack. In the context of *ExtRightCoax*(*st*) we trivially know all the bases required to compute the energy, so we do it there. These ideas of a *GU, WC* and *RightCoax* function are repeated in the *Unpaired* and *Unpaired*2 functions.

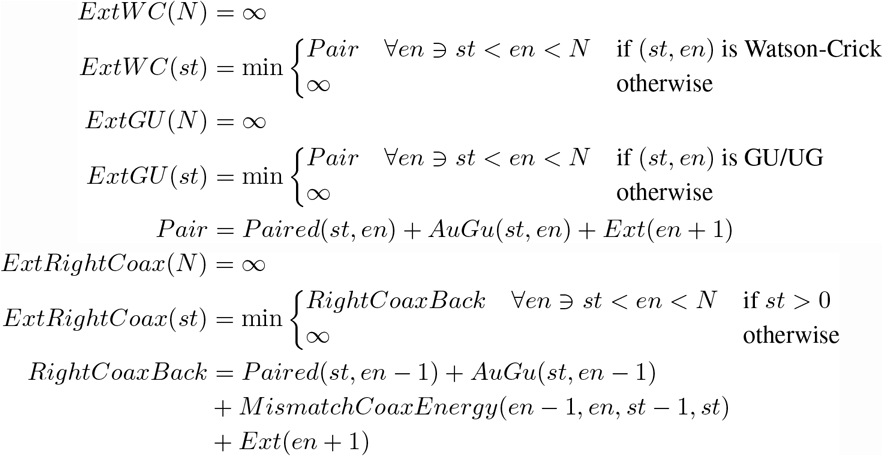

The *Paired*(*st, en*) function returns the energy of the best structure given (*st, en*) is paired. There are many more cases we need to handle, but that is it.

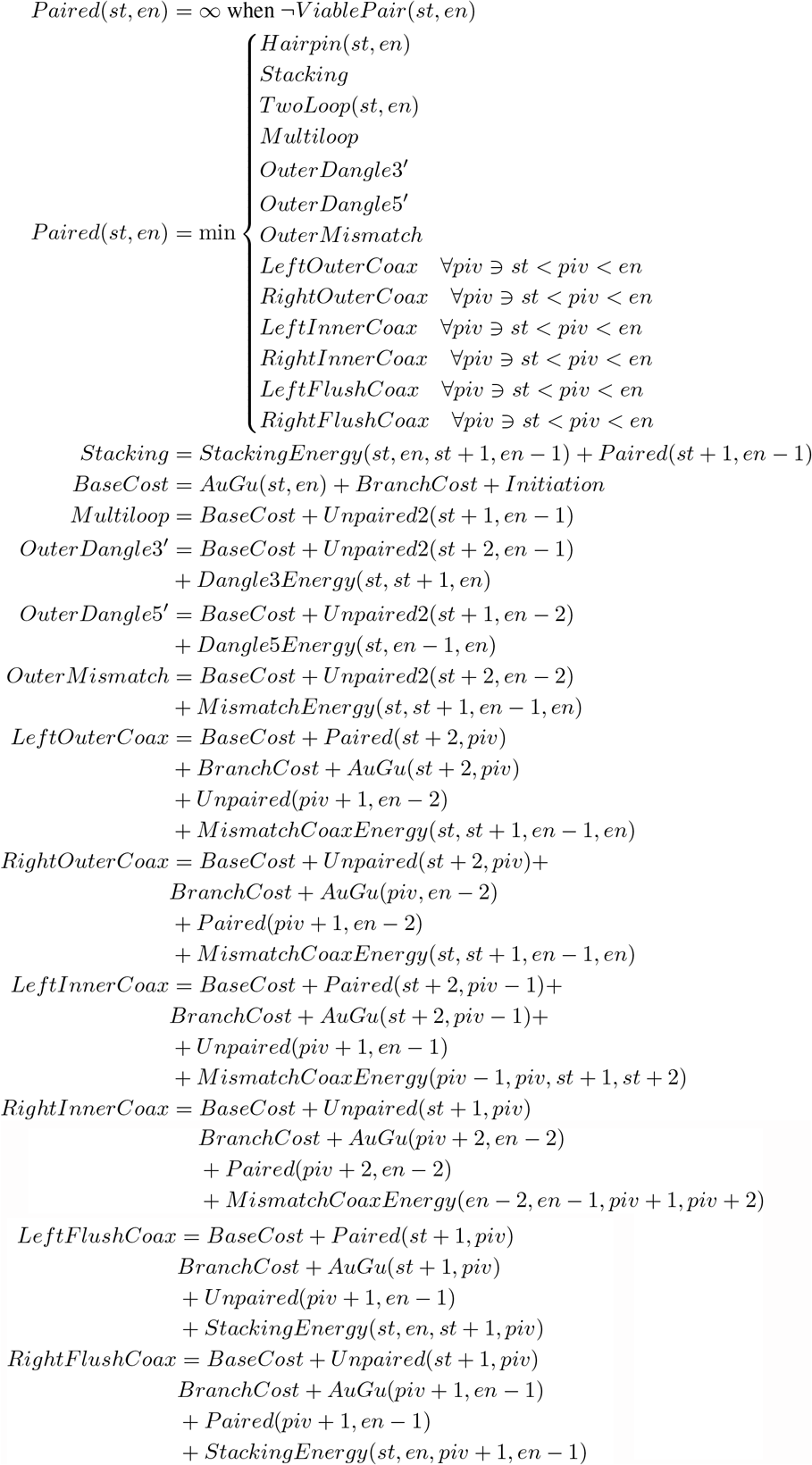

In Table 2, we provide a visual aid for understanding the cases.

The *Unpaired*(*st, en*) function finds the energy of the best structure with at least one branch in the range (*st, en*).

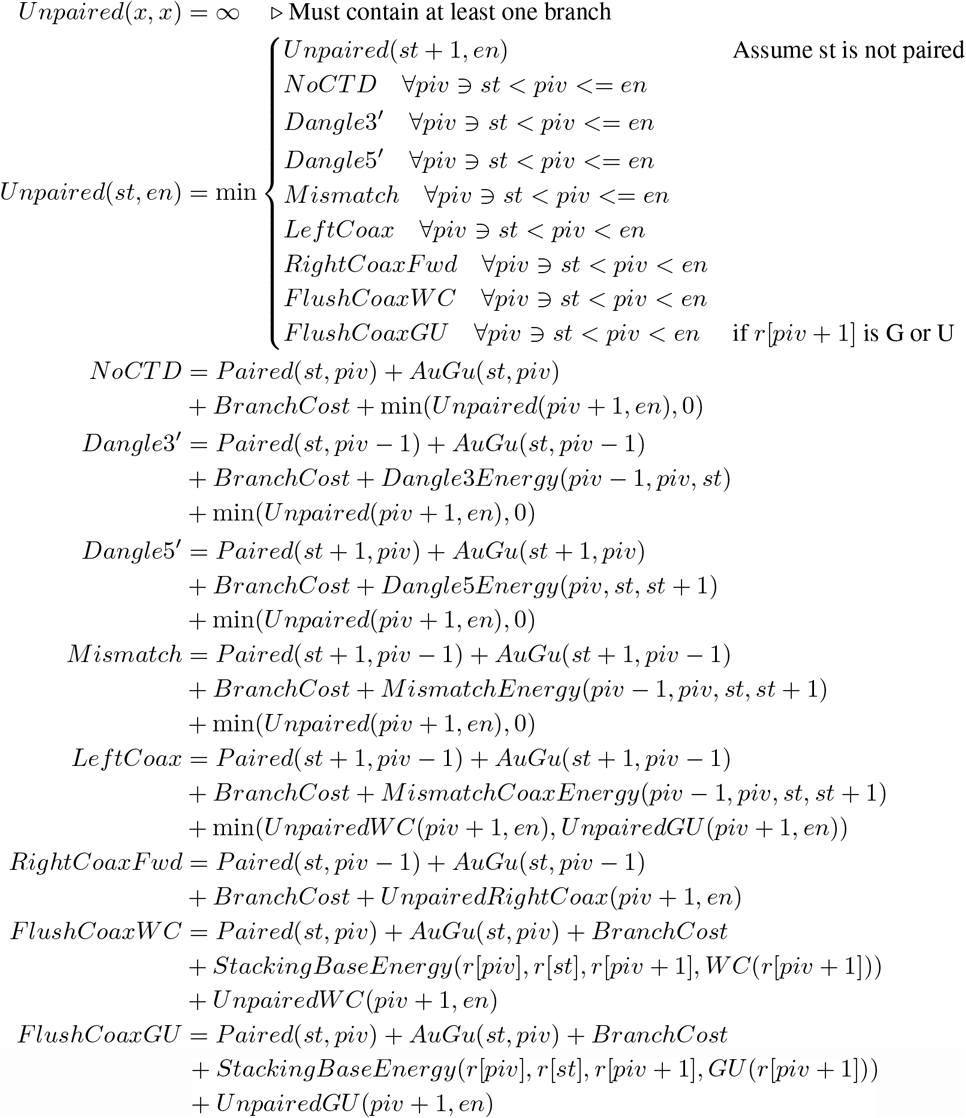

Table 1 shows a visual guide for the cases for *Unpaired* and *Unpaired*2.

The *Unpaired*2 function is mostly the same as *Unpaired*. The min(*Unpaired*, 0) turns into just *Unpaired*, because we need at least two branches.

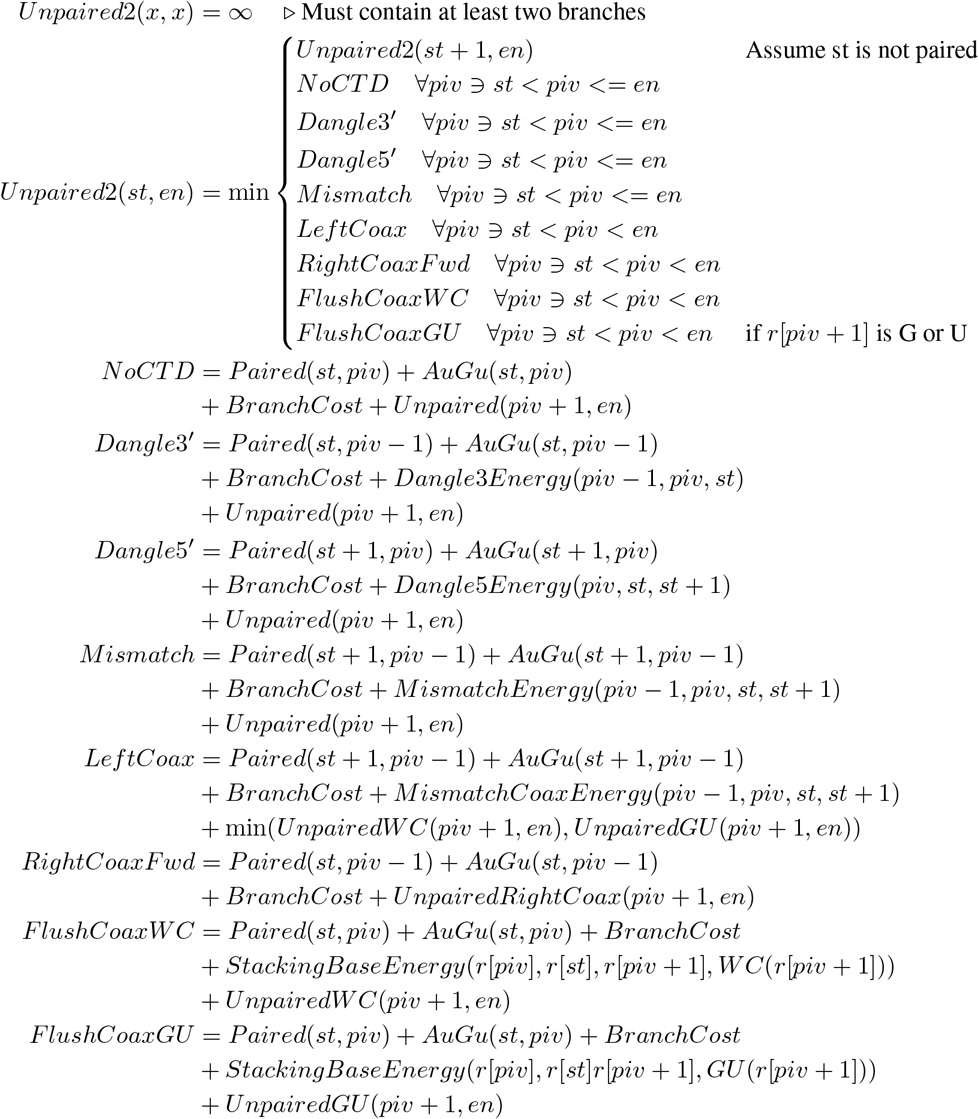

Finally, we have the three functions that compute the helper values for coaxial stacking.

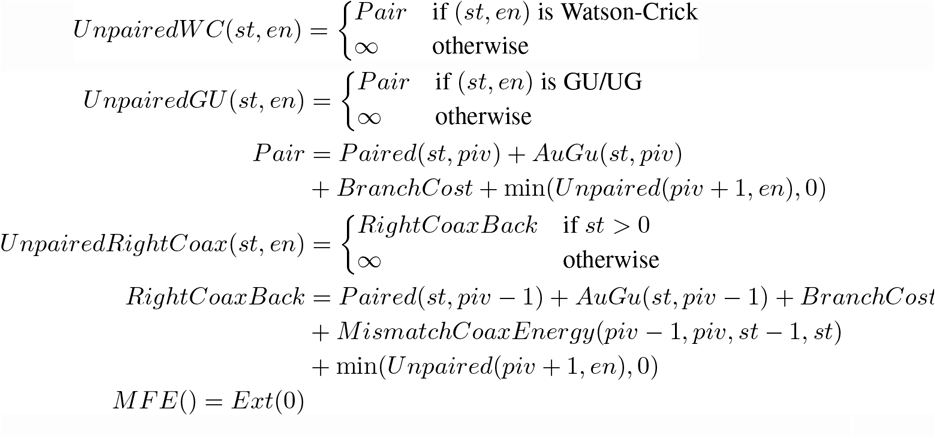

This gives a canonical formulation of the Zuker-Stiegler algorithm that does not need to look at all combinations of two branches in a range to do coaxial stacking. This enables us to apply sparse folding.

## Footnotes

1 This Ninio maximum asymmetry term is not documented in the NNDB anywhere, but it is implemented by RNAstructure, and is alluded to by Lyngsø [59]

## References

[1] Y. Kodama, M. Shumway, and R. Leinonen, “The sequence read archive: Explosive growth of sequencing data,” Nucleic Acids Research, vol. 40, no. D1, pp. D54–D56, Oct. 2011.

[2] J. E. Wilusz, H. Sunwoo, and D. L. Spector, “Long noncoding rnas: Functional surprises from the RNA world,” Genes & Development, vol. 23, no. 13, pp. 1494–1504, 2009.

[3] R. Sah, A. J. Rodriguez-Morales, R. Jha, et al., “Complete genome sequence of a 2019 novel coronavirus (sars-cov-2) strain isolated in nepal,” Microbiology Resource Announcements, vol. 9, no. 11, e00169–20, 2020.

[4] P. P. Amaral, M. E. Dinger, T. R. Mercer, and J. S. Mattick, “The eukaryotic genome as an RNA machine,” Science, vol. 319, no. 5871, pp. 1787–1789, 2008.

[5] P. Nissen, J. Hansen, N. Ban, P. B. Moore, and T. A. Steitz, “The structural basis of ribosome activity in peptide bond synthesis,” Science, vol. 289, no. 5481, pp. 920–930, 2000.

[6] J. A. Doudna and T. R. Cech, “The chemical repertoire of natural ribozymes,” Nature, vol. 418, no. 6894, pp. 222–228, Jul. 2002.

[7] I. Tinoco and C. Bustamante, “How RNA folds,” Journal of Molecular Biology, vol. 293, no. 2, pp. 271–281, 1999.

[8] S. Neidle, Principles of nucleic acid structure. Academic Press, 2010.

[9] N. Pace, B. Thomas, and C. Woese, The RNA world, second edition, 1999.

[10] D. Sankoff, “Simultaneous solution of the RNA folding, alignment and protosequence problems,” SIAM Journal on Applied Mathematics, vol. 45, no. 5, pp. 810–825, 1985.

[11] K. Asai and M. Hamada, “RNA structural alignments, part II: Non-sankoff approaches for structural alignments,” in Methods in Molecular Biology, Humana Press, 2014, pp. 291–301.

[12] J. H. Havgaard and J. Gorodkin, “RNA structural alignments, part i: Sankoff-based approaches for structural alignments,” in Methods in Molecular Biology, Humana Press, Dec. 2013, pp. 275–290.

[13] S. Li, H. Zhang, L. Zhang, et al., “LinearTurboFold: Linear-time global prediction of conserved structures for RNA homologs with applications to sars-cov-2,” Proceedings of the National Academy of Sciences, vol. 118, no. 52, e2116269118, 2021.

[14] H. Zhang, L. Zhang, K. Liu, S. Li, D. H. Mathews, and L. Huang, “Linear-time algorithms for RNA structure prediction,” in Methods in Molecular Biology, Springer US, Jul. 2022, pp. 15–34.

[15] T. Fukunaga and M. Hamada, “LinAliFold and CentroidLinAliFold: Fast RNA consensus secondary structure prediction for aligned sequences using beam search methods,” Bioinformatics Advances, vol. 2, no. 1, vbac078, 2022.

[16] J. H. Havgaard, E. Torarinsson, and J. Gorodkin, “Fast pairwise structural rna alignments by pruning of the dynamical programming matrix,” PLOS Computational Biology, vol. 3, no. 10, pp. 1–13, Oct. 2007.

[17] M. Zuker and P. Stiegler, “Optimal computer folding of large rna sequences using thermodynamics and auxiliary information,” Nucleic Acids Research, vol. 9, no. 1, pp. 133–148, Jan. 1981.

[18] D. H. Mathews, J. Sabina, M. Zuker, and D. H. Turner, “Expanded sequence dependence of thermodynamic parameters improves prediction of RNA secondary structure11edited by i. tinoco,” Journal of Molecular Biology, vol. 288, no. 5, pp. 911–940, 1999.

[19] D. H. Mathews, M. D. Disney, J. L. Childs, S. J. Schroeder, M. Zuker, and D. H. Turner, “Incorporating chemical modification constraints into a dynamic programming algorithm for prediction of rna secondary structure,” Proceedings of the National Academy of Sciences, vol. 101, no. 19, pp. 7287–7292, 2004.

[20] M. Andronescu, A. Condon, D. H. Turner, and D. H. Mathews, “The determination of RNA folding nearest neighbor parameters,” in RNA Sequence, Structure, and Function: Computational and Bioinformatic Methods, J. Gorodkin and W. L. Ruzzo, Eds. Totowa, NJ: Humana Press, 2014, pp. 45–70.

[21] T. Xia, J. J. SantaLucia, M. E. Burkard, et al., “Thermodynamic parameters for an expanded nearest-neighbor model for formation of RNA duplexes with watson-crick base pairs,” Biochemistry, vol. 37, no. 42, pp. 14 719–14 735, 1998, PMID: 9778347.

[22] S. M. Freier, R. Kierzek, J. A. Jaeger, et al., “Improved free-energy parameters for predictions of RNA duplex stability.,” Proceedings of the National Academy of Sciences, vol. 83, no. 24, pp. 9373–9377, 1986.

[23] J. A. Jaeger, D. H. Turner, and M. Zuker, “Improved predictions of secondary structures for RNA.,” Proceedings of the National Academy of Sciences, vol. 86, no. 20, pp. 7706–7710, 1989.

[24] R. Lorenz, S. H. Bernhart, C. Hönerzu Siederdissen, et al., “ViennaRNA package 2.0,” Algorithms for Molecular Biology, vol. 6, pp. 1–14, 2011.

[25] J. S. Reuter and D. H. Mathews, “RNAstructure: Software for RNA secondary structure prediction and analysis,” BMC Bioinformatics, vol. 11, no. 1, pp. 1–9, 2010.

[26] M. Zuker, “Mfold web server for nucleic acid folding and hybridization prediction,” Nucleic Acids Research, vol. 31, no. 13, pp. 3406–3415, Jul. 2003.

[27] N. R. Markham and M. Zuker, “Unafold,” Bioinformatics: Structure, Function and Applications, pp. 3–31, 2008.

[28] L. Huang, H. Zhang, D. Deng, et al., “LinearFold: linear-time approximate RNA folding by 5’-to-3’ dynamic programming and beam search,” Bioinformatics, vol. 35, no. 14, pp. i295–i304, Jul. 2019.

[29] H. Zhang, L. Zhang, D. H. Mathews, and L. Huang, “LinearPartition: Linear-time approximation of rna folding partition function and base-pairing probabilities,” Bioinformatics, vol. 36, no. Supplement_1, pp. i258–i267, Jul. 2020.

[30] Y. Song, “Time and space efficient algorithms for RNA folding with the four-russians technique,” CoRR, vol. abs/1503.05670, 2015.

[31] Y. Frid and D. Gusfield, “A simple, practical and complete o-time algorithm for RNA folding using the four-russians speedup,” Algorithms for Molecular Biology, vol. 5, no. 1, p. 13, Jan. 2010.

[32] B. Venkatachalam, D. Gusfield, and Y. Frid, “Faster algorithms for RNA-folding using the four-russians method,” Algorithms for Molecular Biology, vol. 9, no. 1, p. 5, 2014.

[33] R. Nussinov and A. B. Jacobson, “Fast algorithm for predicting the secondary structure of single-stranded RNA.,” Proceedings of the National Academy of Sciences, vol. 77, no. 11, pp. 6309–6313, 1980.

[34] R. Backofen, D. Tsur, S. Zakov, and M. Ziv-Ukelson, “Sparse RNA folding: Time and space efficient algorithms,” Journal of Discrete Algorithms, vol. 9, no. 1, pp. 12–31, 2011, 20th Anniversary Edition of the Annual Symposium on Combinatorial Pattern Matching (CPM 2009).

[35] S. Will and H. Jabbari, “Sparse RNA folding revisited: Space-efficient minimum free energy prediction,” in Algorithms in Bioinformatics, Springer, 2015, pp. 257–270.

[36] A. E. Walter, D. H. Turner, J. Kim, et al., “Coaxial stacking of helixes enhances binding of oligoribonucleotides and improves predictions of RNA folding.,” Proceedings of the National Academy of Sciences, vol. 91, no. 20, pp. 9218–9222, 1994.

[37] E. Courtney, Edgeworth/memerna: V0.1, version 0.1, 10.5281/zenodo.8214642,https://github.com/Edgeworth/memerna/tree/release/0.1, 2023.

[38] I. Tinoco, O. C. Uhlenbeck, and M. D. Levine, “Estimation of secondary structure in ribonucleic acids,” Nature, vol. 230, no. 5293, pp. 362–367, 1971.

[39] I. Tinoco, P. N. Borer, B. Dengler, et al., “Improved estimation of secondary structure in ribonucleic acids,” Nature, vol. 246, no. 150, pp. 40–41, 1973.

[40] D. H. Turner and D. H. Mathews, “NNDB: the nearest neighbor parameter database for predicting stability of nucleic acid secondary structure,” Nucleic Acids Research, vol. 38, no. suppl_1, pp. D280–D282, Oct. 2009.

[41] D. H. Mathews and D. H. Turner, “Experimentally derived nearest-neighbor parameters for the stability of RNA three- and four-way multibranch loops,” Biochemistry, vol. 41, no. 3, pp. 869–880, 2002, PMID: 11790109.

[42] W. Salser, “Globin mRNA sequences: Analysis of base pairing and evolutionary implications,” Cold Spring Harbor Symposia on Quantitative Biology, vol. 42, pp. 985–1002, 1978.

[43] I. L. Hofacker, W. Fontana, P. F. Stadler, L. S. Bonhoeffer, M. Tacker, P. Schuster, et al., “Fast folding and comparison of RNA secondary structures,” Monatshefte fur chemie, vol. 125, pp. 167–167, 1994.

[44] M. Zuker and D. Sankoff, “RNA secondary structures and their prediction,” Bulletin of mathematical biology, vol. 46, no. 4, pp. 591–621, 1984.

[45] J. Zuber, S. J. Schroeder, H. Sun, D. H. Turner, and D. H. Mathews, “Nearest neighbor rules for RNA helix folding thermodynamics: improved end effects,” Nucleic Acids Research, vol. 50, no. 9, pp. 5251–5262, May 2022.

[46] M. Ward, A. Datta, M. Wise, and D. H. Mathews, “Advanced multi-loop algorithms for RNA secondary structure prediction reveal that the simplest model is best,” Nucleic Acids Research, vol. 45, no. 14, pp. 8541–8550, Jun. 2017.

[47] D. Sankoff and J. B. Kruskal, “Time warps, string edits, and macromolecules: The theory and practice of sequence comparison,” Reading: Addison-Wesley Publication, 1983, edited by Sankoff, David; Kruskal, Joseph B., vol. 1, 1983.

[48] S. Mainville, “Comparaisons et auto-comparaisons de chaînes finies,” Ph.D. dissertation, University of Montreal, Canada, 1981.

[49] D. H. Mathews and D. H. Turner, “Experimentally derived nearest-neighbor parameters for the stability of RNA three- and four-way multibranch loops,” Biochemistry, vol. 41, no. 3, pp. 869–880, Dec. 2001.

[50] J. M. Diamond, D. H. Turner, and D. H. Mathews, “Thermodynamics of three-way multibranch loops in RNA,” Biochemistry, vol. 40, no. 23, pp. 6971–6981, May 2001.

[51] B. Liu, J. M. Diamond, D. H. Mathews, and D. H. Turner, “Fluorescence competition and optical melting measurements of RNA three-way multibranch loops provide a revised model for thermodynamic parameters,” Biochemistry, vol. 50, no. 5, pp. 640–653, Jan. 2011.

[52] Y. Wexler, C. Zilberstein, and M. Ziv-Ukelson, “A study of accessible motifs and RNA folding complexity,” in Research in Computational Molecular Biology, Springer Berlin Heidelberg, 2006, pp. 473–487.

[53] S. Wuchty, W. Fontana, I. L. Hofacker, and P. Schuster, “Complete suboptimal folding of RNA and the stability of secondary structures,” Biopolymers, vol. 49, no. 2, pp. 145–165, 1999.

[54] RNAFOLD, http://www.tbi.univie.ac.at/RNA/RNAfold.1.html, Accessed: 2022/08/22.

[55] P. Virtanen, R. Gommers, T. E. Oliphant, et al., “SciPy 1.0: Fundamental algorithms for scientific computing in python,” Nature Methods, vol. 17, pp. 261–272, 2020.

[56] M. L. Waskom, “Seaborn: Statistical data visualization,” Journal of Open Source Software, vol. 6, no. 60, p. 3021, 2021.

[57] S. Seabold and J. Perktold, “Statsmodels: Econometric and statistical modeling with python,” in 9th Python in Science Conference, 2010.

[58] E. Kierzek, X. Zhang, R. M. Watson, et al., “Secondary structure prediction for rna sequences including n6-methyladenosine,” Nature Communications, vol. 13, no. 1, p. 1271, 2022.

[59] R. B. Lyngsø, M. Zuker, and C. N. S. Pedersen, “An improved algorithm for RNA secondary structure prediction,” BRICS Report Series, vol. 6, no. 15, Jan. 1999.

